# Broiler house environment and litter management practices impose selective pressures on antimicrobial resistance genes and virulence factors of *Campylobacter*

**DOI:** 10.1101/2023.02.02.526821

**Authors:** Reed Woyda, Adelumola Oladeinde, Dinku Endale, Timothy Strickland, Jodie Plumblee Lawrence, Zaid Abdo

## Abstract

*Campylobacter* infections are a leading cause of bacterial diarrhea in humans globally. Infections are due to consumption of contaminated food products and are highly associated with chicken meat, with chickens being an important reservoir for *Campylobacter*. Here, we characterized the genetic diversity of *Campylobacter* species detected in broiler chicken litter over three consecutive flocks and determined their antimicrobial resistance and virulence factor profiles. Antimicrobial susceptibility testing and whole genome sequencing were performed on *Campylobacter jejuni* (n = 39) and *Campylobacter coli* (n = 5) isolates. All *C. jejuni* isolates were susceptible to all antibiotics tested while *C. coli* (n =4) were resistant to only tetracycline and harbored the tetracycline-resistant ribosomal protection protein (TetO). Virulence factors differed within and across grow houses but were explained by the isolates’ flock cohort, species and multilocus sequence type. Virulence factors involved in the ability to invade and colonize host tissues and evade host defenses were absent from flock cohort 3 *C. jejuni* isolates as compared to flock 1 and 2 isolates. Our results show that virulence factors and antimicrobial resistance genes differed by the isolates’ multilocus sequence type and by the flock cohort they were present in. These data suggest that the house environment and litter management practices performed imposed selective pressures on antimicrobial resistance genes and virulence factors. In particular, the absence of key virulence factors within the final flock cohort 3 isolates suggests litter reuse selected for *Campylobacter* strains that are less likely to colonize the chicken host.

**Importance:** *Campylobacter* is a leading cause of foodborne illness in the United States due to the consumption of contaminated food products or from mishandling of food products, often associated with chicken meat. *Campylobacter* is common in the microbiota of avian and mammalian gut; however, the acquisition of antimicrobial resistance genes and virulence factors may result in strains that pose a significant threat to public health. Although there are studies that have investigated the genetic diversity of *Campylobacter* strains isolated from post-harvest chicken samples, there is limited data on the genome characteristics of isolates recovered from pre-harvest broiler production. In this study, we show that *Campylobacter jejuni* and *Campylobacter coli* that differ in their carriage of antimicrobial resistance and virulence factors may differ in their ability to evade host defense mechanisms and colonize the gut of chickens and humans. Furthermore, we found that differences in virulence factor profiles were explained by the species of *Campylobacter* and its multilocus sequence type.

## Introduction

Campylobacteriosis is a leading cause of diarrheal illness worldwide and poultry are the major reservoir of *Campylobacter* species (Young, Davis and DiRita, 2007; De Vries *et al*., 2018). The Centers for Disease Control and Prevention (CDC, 2019) estimate that 1.5 million United States residents are affected by campylobacteriosis each year (CDC, 2019). While it rarely results in long-term health problems, studies estimate that 5-20% of campylobacteriosis cases develop irritable bowel syndrome, 1-5% develop arthritis and, in very rare instances, campylobacteriosis may cause Guillain-Barré syndrome (Mishu 1993, Hansson 2016). Economic burden from *Campylobacter* infections was estimated in 2012 to be $1.56 billion and *Campylobact*er, specifically from poultry, wa*s* ranked as the leading pathogen-food combination to cause health risks to humans and to negatively impact the economy (Scharff, 2012; Hoffmann, Maculloch and Batz, 2015). Transmission of *Campylobacter* species occurs through consumption or handling of contaminated food products, direct contact with farm or domesticated animals and abattoir workers not practicing good handwashing and food safety practices (Hansson *et al*., 2018; Igwaran and Okoh, 2019; Mourkas *et al*., 2020).

*Campylobacter* pathogenicity, disease severity and treatment options are influenced by the repertoire of virulence factors (VFs) and antimicrobial resistance genes (ARGs) they carry. The global rise of antimicrobial resistance (AMR) has impaired effective treatment of *Campylobacter* infections especially when *Campylobacter* strains harbor ARGs that confer resistance to critically important antibiotics (Florez-Cuadrado *et al*., 2016; Chen *et al*., 2018; Liu *et al*., 2019; Zachariah *et al*., 2021; Liao *et al*., 2022). *Campylobacter* ARGs may be acquired through mutations (for example the C257T change in *gyrA* (DNA gyrase)) resulting in resistance to fluoroquinolones), encoded on plasmids, or located within multidrug resistance genomic islands (MDRGIs) such as the *erm*(B), which confers high levels of macrolide resistance (Shen *et al*., 2018).

Importantly, ARGs located on plasmids or MDRGIs are generally transferable across *Campylobacter* species which may lead to the emergence of multidrug resistant strains. Similarly, *Campylobacter* may carry VFs that increase their pathogenicity and the ability to survive within a given host, which can exacerbate disease severity (Lopes *et al*., 2021). Like ARGs, VFs may be accumulated in Campylobacters leading to strains that are highly virulent and pathogenic (Ghatak *et al*., 2017; de Fátima Rauber Würfel *et al*., 2020; Lopes *et al*., 2021). This may lead to their persistence through pre-harvest and post-harvest, thereby posing a risk to the public (Ghatak *et al*., 2017; Al Hakeem *et al*., 2022). Furthermore, strains carrying ARGs conferring resistance to critically important antibiotics for humans, as well as possessing VFs that increase their ability to colonize host tissues, will be harder to treat in the event contaminated food products reach consumers (Montgomery *et al*., 2018; Liu *et al*., 2019; Béjaoui *et al*., 2022).

Bacterial pathogens such as *Campylobacter* have been shown to persist in poultry litter that is reused to grow multiple flocks of broiler chickens (Rauber Würfel *et al*., 2019). The copraphagic nature of these birds makes the litter one of the first broiler sourced material ingested upon placement in a broiler house. Therefore, it is plausible that broiler chicks get exposed to *Campylobacter* during pecking, bathing, or resting activities. Many studies have characterized *Campylobacter* presence in commonly used bedding materials such as pine shavings, sawdust and ricehulls (Kelley *et al*., 1995; Pope and Cherry, 2000; Willis, Murray and Talbott, 2000; Stern *et al*., 2001; Line, 2002, p. 2; Line and Bailey, 2006; Kassem *et al*., 2010; Rauber Würfel *et al*., 2019). However, none of these studies performed an in-depth genomic characterization of the *Campylobacter* isolates found.

We previously showed that *C. coli and C. jejuni* were unequally distributed across the litter of four co-located broiler houses on a single farm (Oladeinde *et al*., 2022). We also showed that the probability of detecting *Campylobacter* in litter was higher for the first broiler flock cohort raised on litter compared to cohorts 2 and 3 (Oladeinde *et al*., 2022). In the present study, we performed antimicrobial susceptibility testing and whole genome sequencing on forty-four *Campylobacter* isolates recovered from the study (Oladeinde *et al*., 2022). An in-depth genomic characterization and phylogenetic analysis revealed that isolates clustered based on their VFs and suggest that the litter environment exacts a selective pressure on VFs harbored by *Campylobacter* species.

## Results

### *Campylobacter jejuni* and *coli* were present over three broiler flock cohorts

The objective of this study was to understand the changes in ARGs and VFs in *Campylobacter* isolates obtained from litter used to raise multiple flocks of birds. Isolates were obtained from peanut hull litter samples collected from three consecutive flock cohorts within 4 co-located broiler houses. To determine the prevalence of *Campylobacter* in the collected litter samples (Oladeinde *et al*., 2022), we performed direct and selective enrichment plating of litter eluate onto Cefex agar (Oladeinde *et al*., 2022). *Campylobacter* was detected in 9.38 % (27/288) of litter samples. Next, we selected a total of 44 *Campylobacter* isolates for whole genome sequencing (Oladeinde *et al*., 2022). At least one Campylobacter isolate was selected from each Campylobacter positive litter sample. Additionally, if there were multiple positive isolates obtained from the same litter sample using the different isolation methods (i.e., direct plating, filter method, or enrichment), the filter and enrichment isolates were chosen over the direct plating isolates. The sequenced *Campylobacter* population consisted of 5 *Campylobacter coli* (CC) and 39 *Campylobacter jejuni* (CJ) isolates as determined by taxonomic classification via Kraken2 (Wood, Lu and Langmead, 2019). *C. coli* was isolated from flocks 1 (n=1) and 2 (n=4) samples but absent from flock 3 (**Table 1**), while *C. jejuni* was isolated from each of the 3 flock cohorts. Occurrence of *C. jejuni* across flock cohorts was predominantly in flocks 1 (n=30) and 3 (n=8), while only 1 *C. jejuni* was isolated in flock 2. The 4 co-located broiler houses had unequal representation of both *C. coli* and *C. jejuni* (**Figure 1**, (Oladeinde *et al*., 2022)). Houses 1, 2, 3 and 4 harbored 3, 3, 17 and 16 *C. jejuni* isolates respectively, while houses 1, 3 and 4 harbored 1, 3, and 1 *C. coli* isolates, respectively. Only broiler house 4 harbored *C. jejuni* over 3 consecutive flock cohorts and no house harbored *C. coli* over successive flocks. All *C. jejuni* isolated during flocks 1 and 2 were ST464 while all flock 3 isolates were ST48 (**Table S1**). One *C. coli* isolate was identified as ST9450, while the remaining 4 *C. coli* isolates did not match any multilocus sequence type (MLST) profiles (**Table S1**) due to poor sequencing coverage. Taken together, *C. jejuni* and *C. coli* were isolated from peanut hull-based litter reused to raise 3 consecutive flocks of birds within 4 co-located broiler houses.

**Figure 1.**
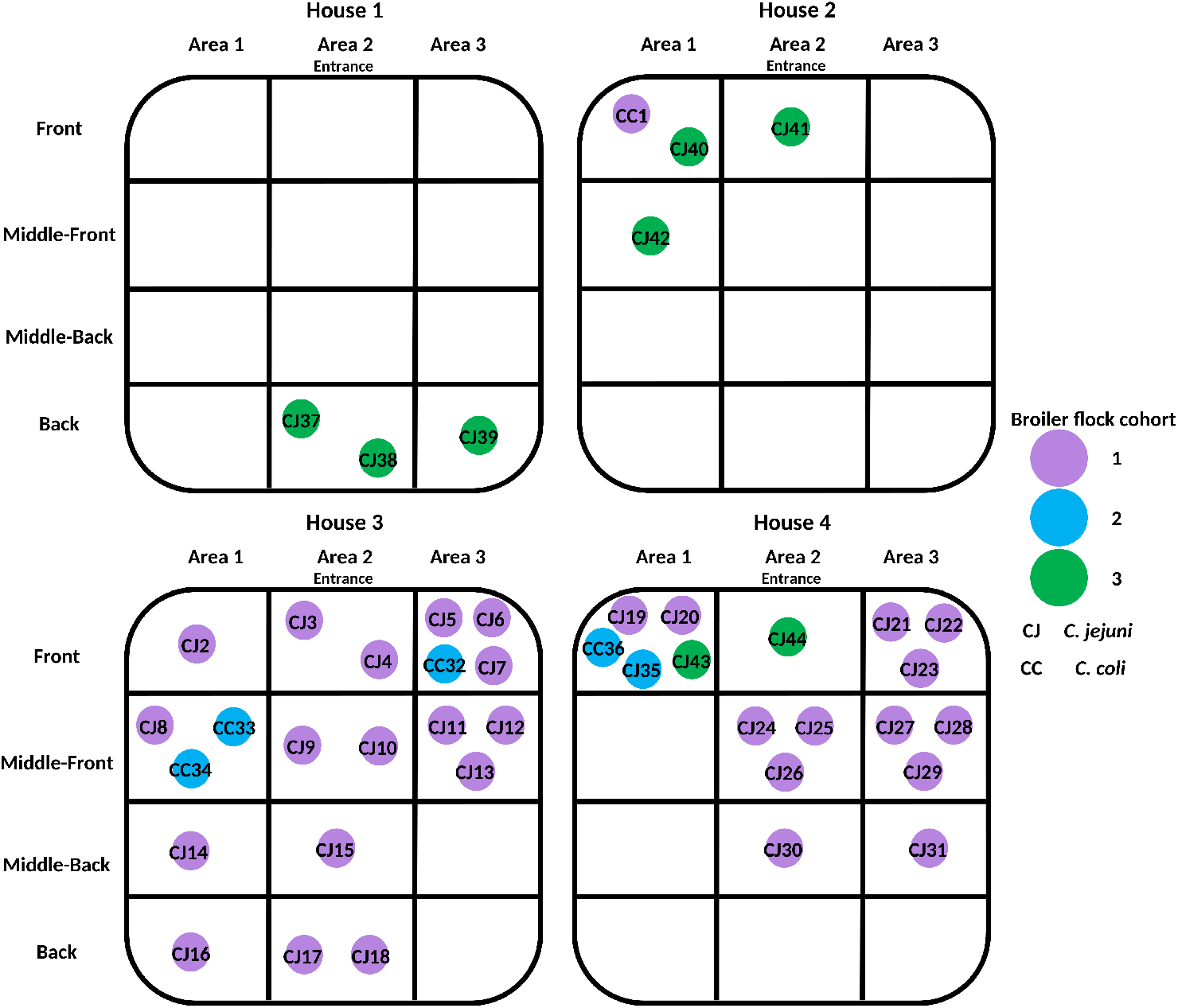
Visual representation of *Campylobacter* species isolated from peanut hull litter within each section of each broiler house. Samples were taken from 4 co-located broiler houses from 3 consecutive flock cohorts. Circles represent individual isolates labeled by their prospective species: *Campylobacter jejuni* (CJ) and *Campylobacter coli* (CC). Circle color indicates which flock cohort an isolate was obtained from: flock cohort 1 (purple), flock cohort 2 (blue) and flock cohort 3 (green).

**Table 1.** Occurrence of Campylobacter species in peanut hull litter from 3 consecutive grow-out cycles across 4 co-located broiler houses.

| Broiler<br>House | Flock cohort 1 |  | Flock cohort 2 |  | Flock cohort 3 |  |
| --- | --- | --- | --- | --- | --- | --- |
|  | *C. coli* | *C. jejuni* | *C. coli* | *C. jejuni* | *C. coli* | *C. jejuni* |
| 1 | 0 | 0 | 0 | 0 | 0 | 3 |
| 2 | 1 | 0 | 0 | 0 | 0 | 3 |
| 3 | 0 | 17 | 3 | 0 | 0 | 0 |
| 4 | 0 | 13 | 1 | 1 | 0 | 2 |

### Virulence factor and antimicrobial resistance profiles differed by species and by multilocus sequence type

We sought to understand the relationship between an isolate’s repertoire of ARGs and VFs, its spatial distribution within a given broiler house, and how it is impacted by broiler house environmental factors. Environmental parameters measured included house temperature, litter moisture, and litter pH. Hierarchical clustering based on the presence and absence of all identified VFs and ARGs (**Table S1**) revealed ARG and VF profiles were grouped by flock, *Campylobacter* species and by isolates’ multilocus sequence type (**Figure 2**). Correspondence analysis revealed overlapping 95% confidence ellipses for isolates by species (**Figure 3A**). While hierarchical clustering grouped *C. jejuni* isolates by flock, it is important to note *C. jejuni* isolates from flock 1 encompass a single MLST, ST464, while flock 3 isolates are all ST48. This was recapitulated through correspondence analysis which identified non-overlapping 95% confidence ellipses for isolates by MLST (**Figure 3B**). Taken together, these results suggest that the species and sequence type of *Campylobacter* are the main factors explaining the differences observed in ARG and VF profiles.

**Figure 2.**
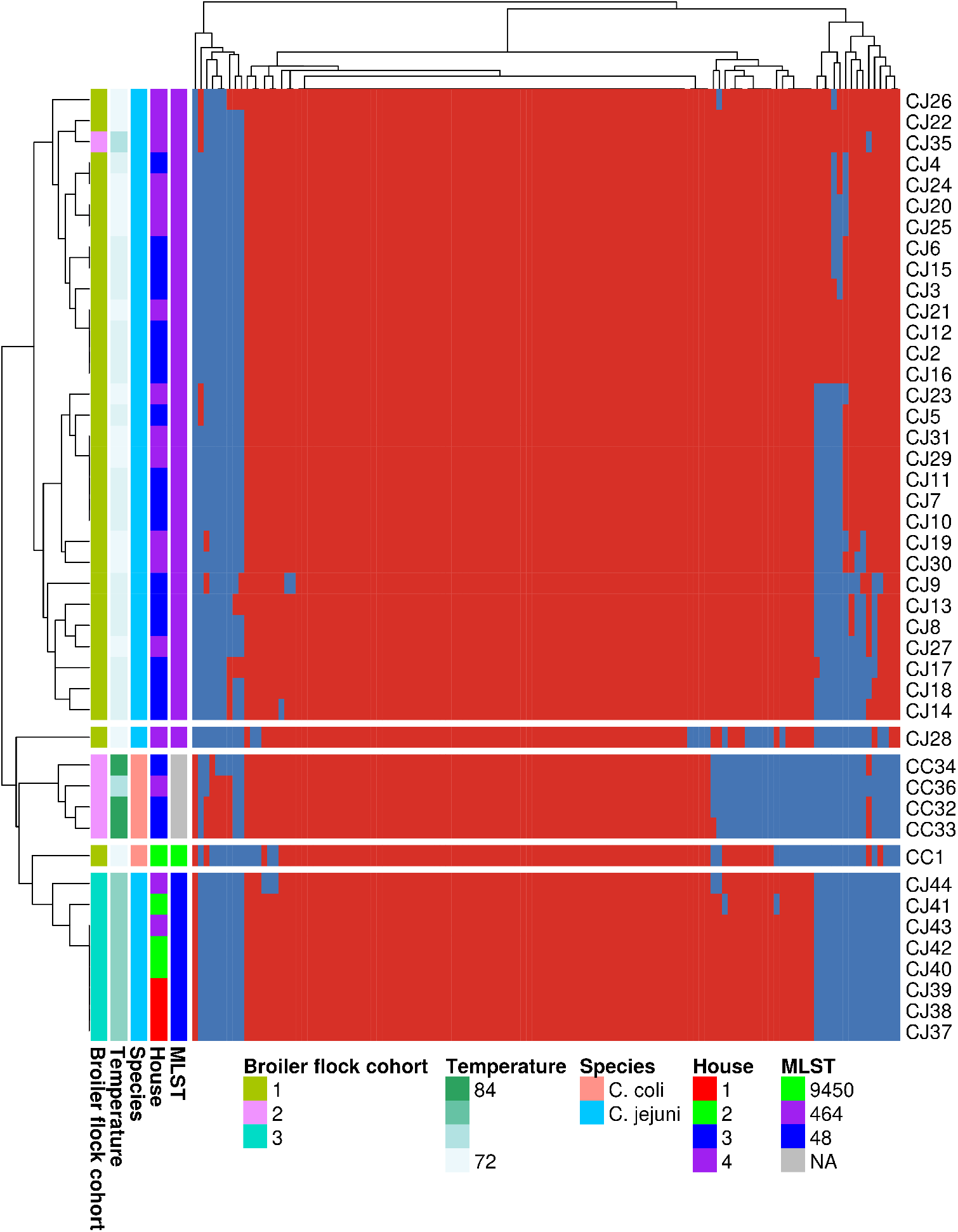
Heatmap of antimicrobial resistance genes and virulence factors across all isolates. Hierarchical clustering revealed ARG and VF profiles were grouped by flock, Campylobacter species and by isolates’ multilocus sequence type. Heatmap was generated in R v4.0.4 with pheatmap v1.0.12 (clustering_method = “average” (UPGMA), clustering_distance_cols = “binary”) using the filtered antimicrobial resistance gene and virulence factor table (**Table S1**). Columns on the right-hand side display the metadata associated with each isolate (labels on left-hand side).

**Figure 3.**
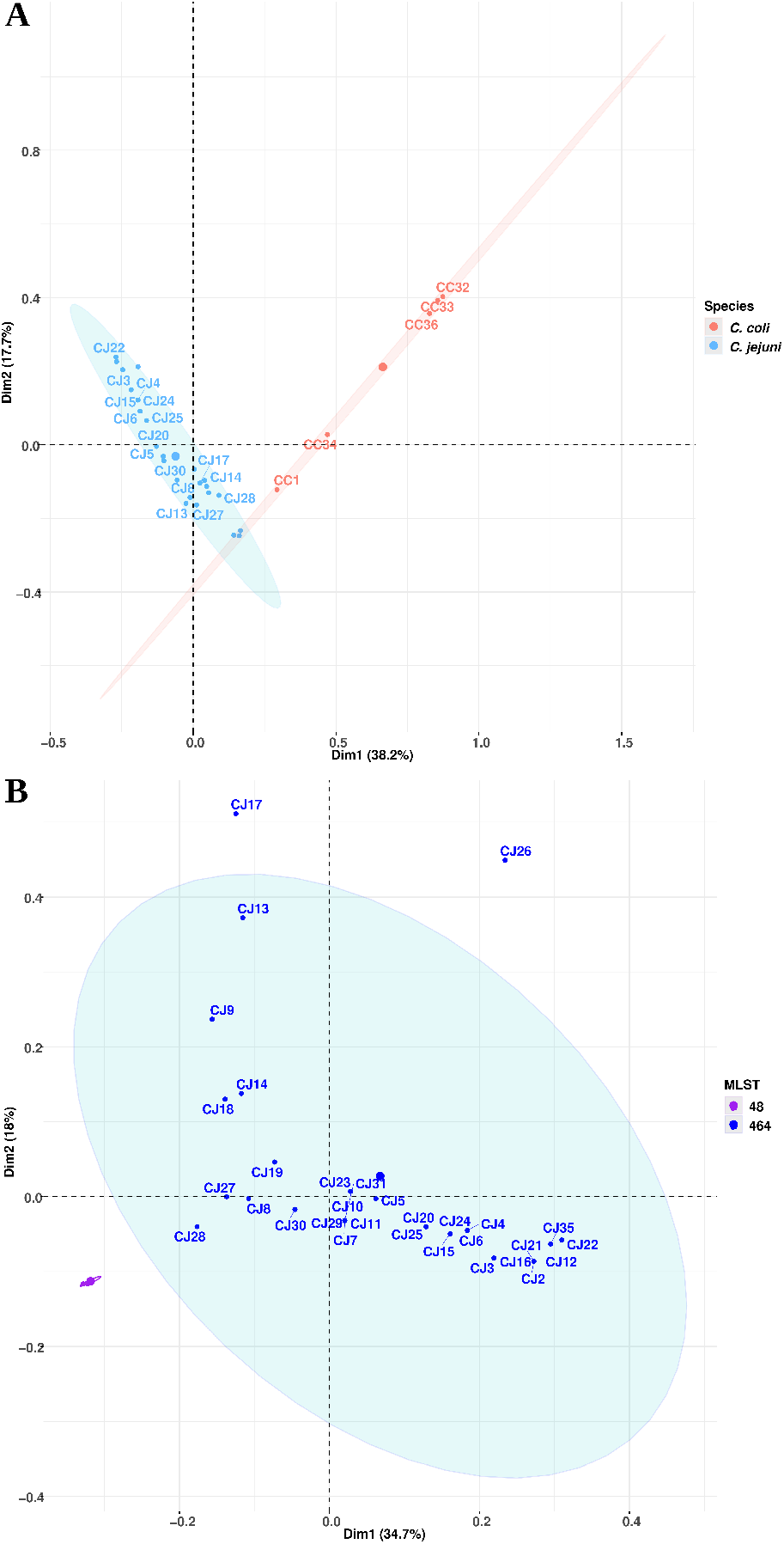
Correspondence analysis using presence/absence ARG and virulence factor profiles. Correspondence analysis revealed overlapping 95% confidence ellipses for isolates by species and non-overlapping 95% confidence ellipses for isolates by MLST. **(A)** Correspondence analysis on all *Campylobacter* isolates (n=44). Circle color corresponds to the verified species: *C. coli* (red) and *C. jejuni* (blue). Species labels are denoted with CC (*C. coli*) and CJ (*C. jejuni*). **(B)** Correspondence analysis on *Campylobacter jejuni* isolates (n = 39). Circle color corresponds to the identified multilocus sequence type (MLST). Ellipses represent a 95% confidence ellipsoid. Correspondence analysis on the presence/absence table of ARGs and virulence factors was conducted in R using factoextra v1.0.7, FactoMineR v2.4 and corrplot v0.2-0 packages.

### Virulence factor profiles differed between species and multilocus sequence type

Correspondence and clustering analysis revealed that VF and ARG differences were explained by species and sequence type. Therefore, we sought to further investigate the functional differences of VFs across these parameters. The 112 VFs identified were grouped into 10 VF functional categories (**Table S1, Table S2**). Sixty-three (56%) of the VFs were present in all isolates (**Table S1**). Across species, *C. jejuni* isolates harbored significantly higher average proportions of VFs than *C. coli* (P< 0.01) with functions relating to toxins, adherence, invasion, colonization and immune evasion, and motility and export apparatus (**Figure 4**, **Table 2**). *C. coli* isolate CC1 was the sole *C. coli* isolate that harbored *Cj1415*/*cysC* (cytidine diphosphoramidate kinase) - a toxin-related VF. *Cj1415*/*cysC* is involved in polysaccharide modification and contributes to serum resistance and the invasion of epithelial cells (Taylor and Raushel, 2018). Contrastingly, *Cj1415*/*cysC* was present in all *C. jejuni* isolates except for isolate 28 (CJ28). The *ctdA*, *ctdB* and *ctdC* VFs encoding for the cytolethal-distending toxin and responsible for cellular distension and death in the epithelial cell layer, were present in all *C. jejuni* isolates. The CheVAWY (*cheA, cheY, cheV* and *cheW*) system that is involved in adherence, motility, and chemotaxis (Reuter *et al*., 2021), was also present in all *C. jejuni* isolates. Moreover, studies have shown that gene deletions, or insertional inactivation of *cheY* can result in the attenuation of growth within the chicken gastrointestinal tract (Hendrixson and DiRita, 2004) and the inability to colonize in murine or ferret disease models (Yao, Burr and Guerry, 1997; Bereswill *et al*., 2011).

**Figure 4.**
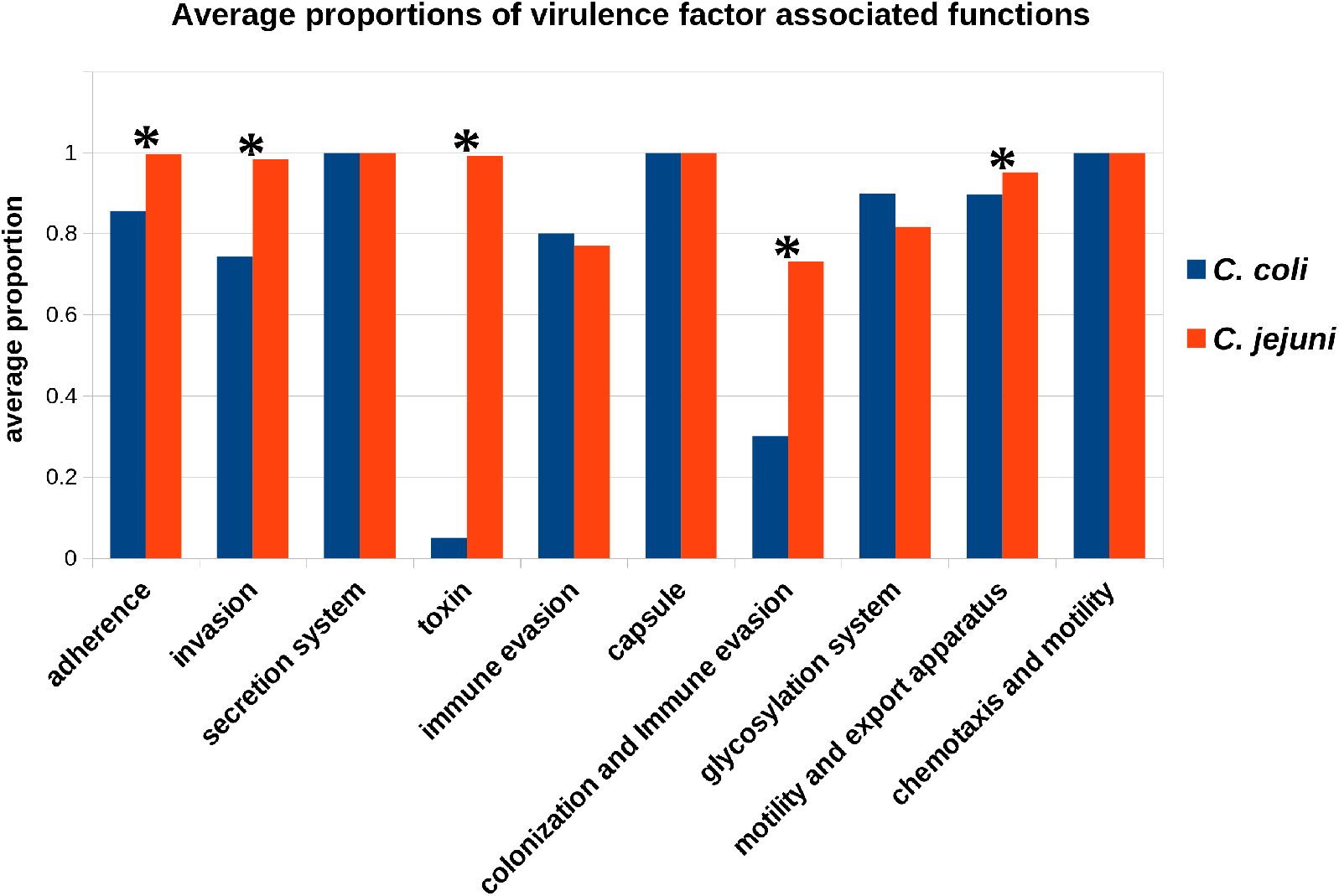
Average proportion of virulence factor-associated functions between *C. coli* and *C. jejuni* isolates. *C. jejuni* isolates harbored significantly higher average proportions of VFs with functions relating to toxins, adherence, invasion, colonization and immune evasion, and motility and export apparatus. Comparison was performed using the Wilcoxon rank-sum test. (**\***) indicates an adjusted p value > 0.05. Virulence genes associated with each function were enumerated for each isolate and a proportion was calculated using the total number of genes in the study population with the given function. Adjusted p value adjustment was performed by the Benjamini-Hochberg false discovery rate correction.

**Table 2.** Average proportion of virulence factor-associated functions between C. coli and C. jejuni isolates. Comparison was performed using the Wilcoxon rank-sum test. Virulence genes associated with each function were enumerated for each isolate and a proportion was calculated using the total number of genes in the study population with the given function. Adjusted p value adjustment was performed using the Benjamini-Hochberg false discovery rate correction method. ‘NaN’ values are due to the inability to compute p values due to average proportion values being identical for all isolates.

| **Virulence Factor Function** | **adjusted p value** |
| --- | --- |
| Toxin | 1.76E-08 |
| Adherence | 4.16E-08 |
| Invasion | 3.01E-06 |
| Motility and export apparatus | 3.44E-06 |
| Colonization and Immune evasion | 0.0062 |
| Glycosylation system | 0.31 |
| Immune evasion | 1 |
| Capsule | NaN |
| Chemotaxis and motility | NaN |
| Secretion system | NaN |

The capsular polysaccharrides (CPS) of *C. jejuni* are involved in virulence and are essential for survival in certain host environments. The CPS transporter gene (*kpsE*) was present in all *C. jejuni* isolates and in *C. coli* isolate 1 (CC1). *C. jejuni* lipopolysaccharide (LPS) is a known VF that mediates adhesion to epithelial cells while *kpsM* and *kpsT* are involved in LPS export (Karlyshev *et al*., 2002). Both *kpsM* and *kpsT* were present in all *C. jejuni* and *C. coli* isolates. However, *kpsC* that is responsible for capsule modification, was only present in *C. jejuni* isolates and in a single *C. coli* isolate, CC1.

When comparing the presence of VFs in *C. jejuni* isolates across houses, only VF functions relating to immune evasion and glycosylation system were significantly different (P<0.01) (**Table 3**, **Table 4**). *C. jejuni* isolates from house 3 harbored more VF functions relating to immune evasion and glycosylation than *C. jejuni* isolates from houses 1 (Table 3) and 2 (Table 4). No significant differences in VF functions were found between house 1 and house 2 *C. jejuni* isolates, or between house 4 and any other house (**Table S3, S4, S5, S6**). When compared across flocks, significant differences in VF functions relating to immune evasion, glycosylation system, and ‘colonization and immune evasion’ were found between *C. jejuni* isolates from flock 1 and flock 3 (**Table 5**). No significant differences in VF functions were found between flock 1 and 2 *C. jejuni* isolates (**Table S7**) albeit, only 1 *C. jejuni* isolate was sequenced from flock 2.

**Table 3.**
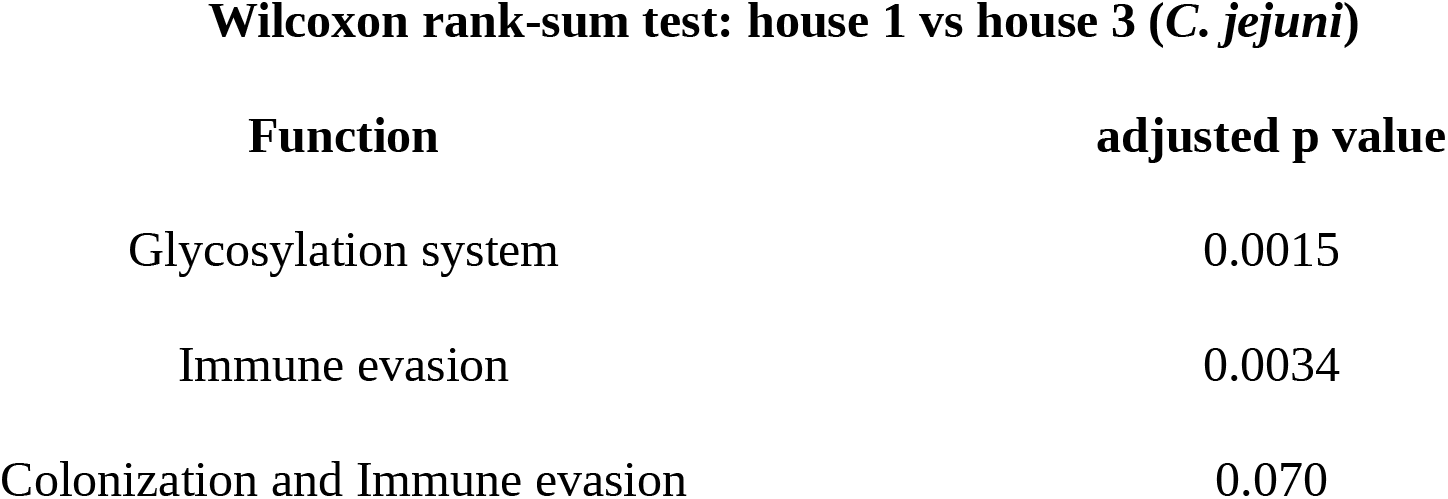
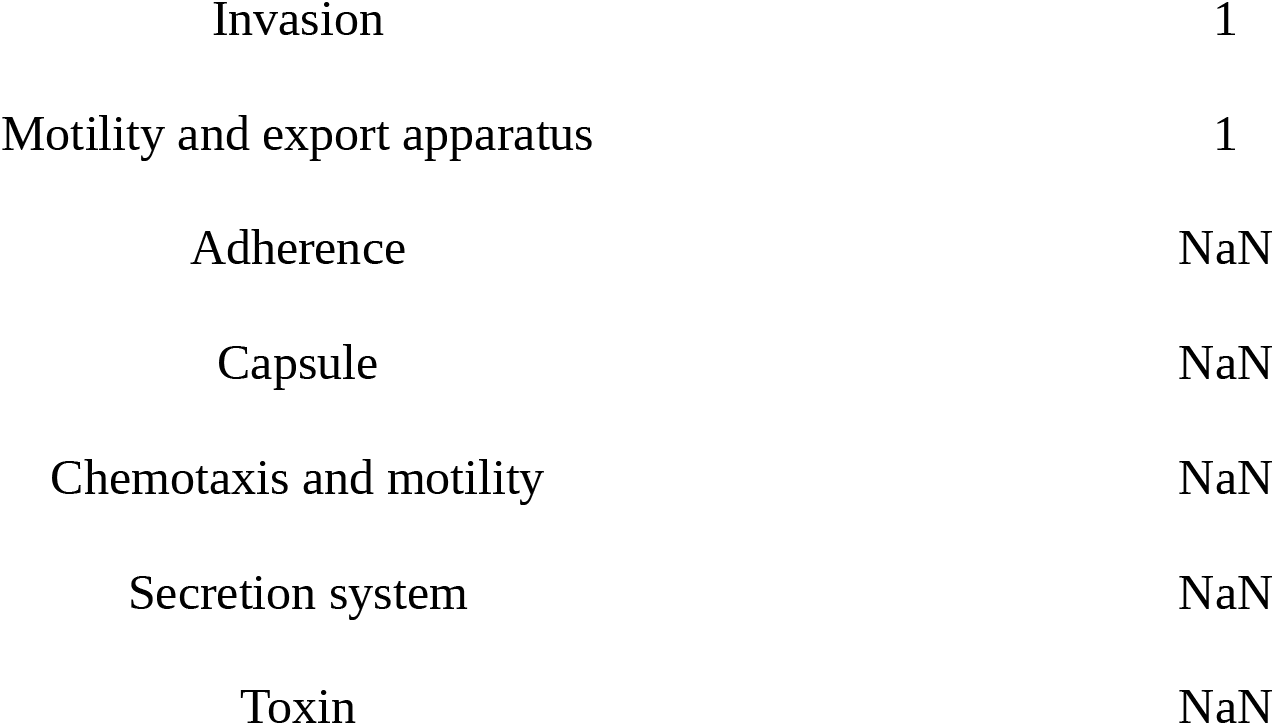
Comparison of average proportions of virulence factor-associated functions for *C. jejuni* isolates which significantly differed in proportion between broiler houses 1 and 3 as determined by the Wilcoxon rank-sum test. Virulence genes associated with each function were enumerated for each isolate and a proportion was calculated using the total number of genes in the study population with the given function. Adjusted p value adjustment was performed by the Benjamini-Hochberg false discovery rate correction method. ‘NaN’ values are due to the inability to compute p values due to average proportion values being identical for all isolates.

**Table 4.** Comparison of average proportions of virulence factor-associated functions for C. jejuni isolates which significantly differed in proportion between broiler houses 2 and 3 as determined by the Wilcoxon rank-sum test. Virulence genes associated with each function were enumerated for each isolate and a proportion was calculated using the total number of genes in the study population with the given function. Adjusted p value adjustment was performed by the Benjamini-Hochberg false discovery rate correction method. ‘NaN’ values are due to the inability to compute p values due to average proportion values being identical for all isolates.

| **Function** | **adjusted p value** |
| --- | --- |
| Glycosylation system | 0.0015 |
| Immune evasion | 0.0034 |
| Colonization and Immune evasion | 0.070 |
| Adherence | 0.25 |
| Invasion | 1 |
| Motility and export apparatus | 1 |
| Capsule | NaN |
| Chemotaxis and motility | NaN |
| Secretion system | NaN |
| Toxin | NaN |

**Table 5.**
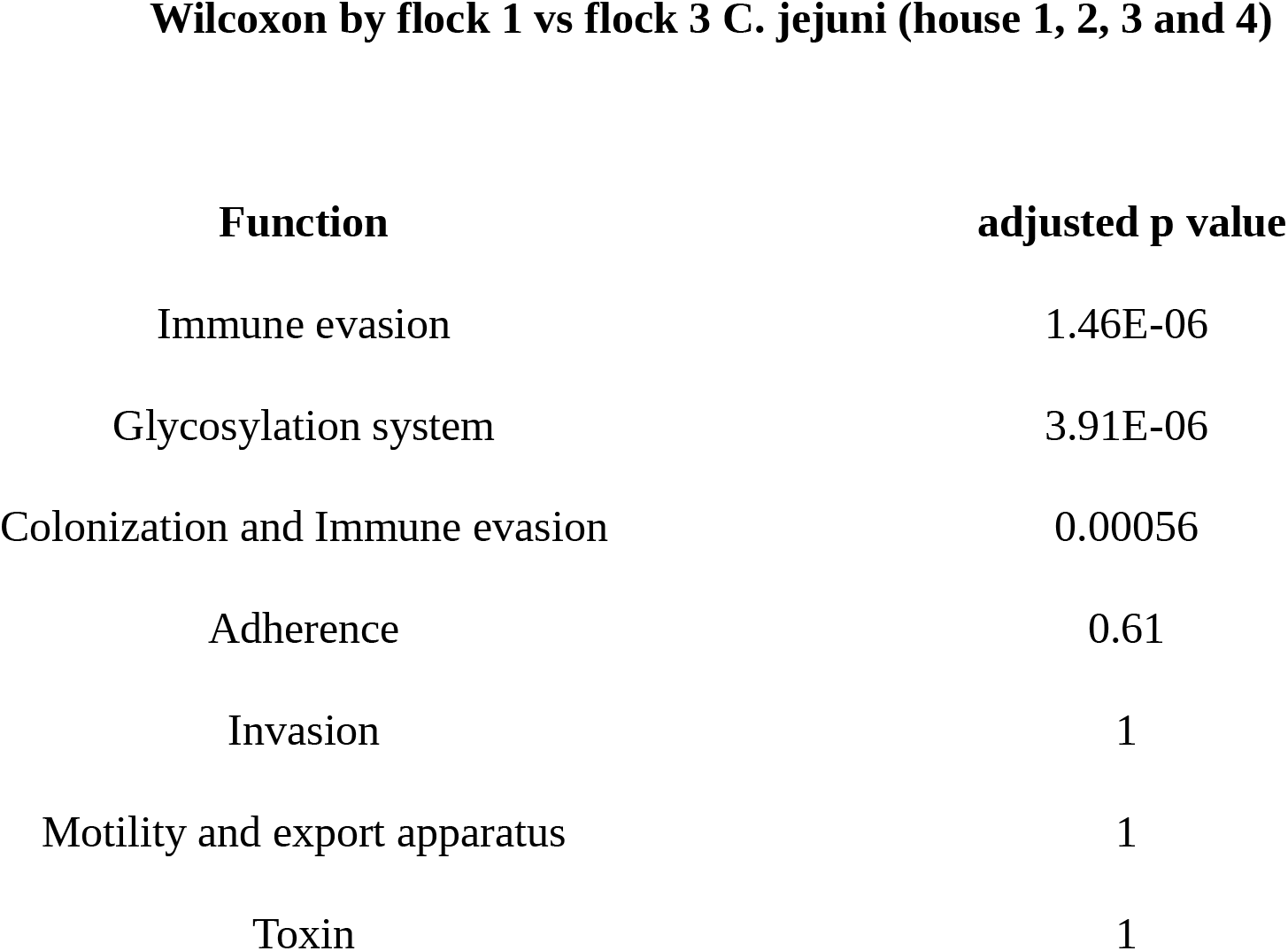
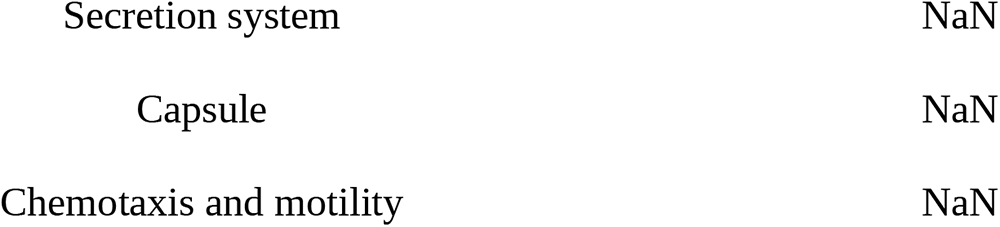
Comparison of average proportions of virulence factor-associated functions for *C. jejuni* isolates which significantly differed in proportion between flock cohorts 1 and 3 as determined by the Wilcoxon rank-sum test. Virulence genes associated with each function were enumerated for each isolate and a proportion was calculated using the total number of genes in the study population with the given function. Adjusted p value adjustment was performed by the Benjamini-Hochberg false discovery rate correction method. ‘NaN’ values are due to the inability to compute p values due to average proportion values being identical for all isolates.

| Function | adjusted p value |
| --- | --- |
| Immune evasion | 1.46E-06 |
| Glycosylation system | 3.91E-06 |
| Colonization and Immune evasion | 0.00056 |
| Adherence | 0.61 |
| Invasion | 1 |
| Motility and export apparatus | 1 |
| Toxin | 1 |
| Secretion system | NaN |
| Capsule | NaN |
| Chemotaxis and motility | NaN |

Similarly, no significant differences in VF were identified between flock 2 and flock 3 *C. jejuni* isolates (**Table S8**). There were 7 VFs (*Cj1426c, fcl, pseE, kfiD, Cj1432c, Cj1440c* and *glf*) that were present in flock 1 (ST464) isolates but absent in *C. jejuni* isolates from flock 3 (ST48) (**Table S9**). *Cj1426c, kfiD, Cj1432c, Cj1440c* and *glf* are all involved in capsule biosysnthesis and transport, *fcl* (putative fucose synthase) is involved in LPS biosynthesis, and *pseE* is involved in O-linked flagellar glycosylation. Taken together, *C. jejuni* isolates from flock 3, all identified as ST48, harbored fewer VF than *C. jejuni* isolates from flocks 1 and 2, identified as ST464.

### The presence of Type IV and type VI secretion systems differentiated *Campylobacter* species

Both type IV and type VI secretion systems (T4SS and T6SS) enable delivery of bacterial effector proteins into neighboring bacterial and eukaryotic cells (Bleumink-Pluym 2013) and are commonly present in *C. jejuni* and *C. coli* (Bleumink-Pluym 2013, Daya Marasini 2020). Although genes encoding these secretion systems may be chromosomally encoded, they are commonly identified on plasmids (Cascales, 2008; Lertpiriyapong *et al*., 2012; Bleumink-Pluym *et al*., 2013; Ghatak *et al*., 2017). Using the *C. jejuni* strain WP2202 plasmid pCJDM202 (NZ_CP014743) that has both the T4SS and T6SS operons as a reference, we determined that T4SS was present only in *C. coli* isolates and T6SS was present only in *C. jejuni* isolates. *C. coli* isolates CC32, CC34 and CC36 harbored the largest segments of the reference query (>88%) while CC33 only contained 33% of the reference query (**Table S10**). Sixteen *C. jejuni* isolates (41%) harbored >85% of the T6SS operon with >99% pairwise identity (**Table S11**). Eight *C. jejuni* isolates contained between 14% and 76% of the T6SS operon, with >98% pairwise identity, while the remaining 15 isolates harbored <10% of the operon. No isolates from the final flock cohort, flock 3, harbored any T6SS genes. In summary, T4SS and T6SS were *Campylobacter* species-specific and were only found in isolates recovered from the litter of flocks 1 and 2. As these secretion systems aid in colonization in chickens, as well as humans, this data suggests isolates that lack these systems may have a decreased ability to infect chickens in subsequent flocks.

### ARG carriage differed by species and flock

Antimicrobial susceptibility testing (AST) was performed on all *Campylobacter* isolates (**Table S12**). All *C. jejuni* isolates were susceptible to all drugs tested, while *C. coli* isolates (4/5) were resistant to tetracycline. Eleven ARGs were identified across all isolates after gene annotation (**Table S13**). All *C. coli* isolates with phenotypic resistance to tetracycline harbored *tetO* (tetracycline resistance gene). *tetO* is known to confer tetracycline resistance in *C. coli* (Sougakoff *et al*., 1987). To determine if *tetO* was harbored on a plasmid, we used *C. jejuni* strain WP2202 plasmid pCJDM202 (NZ_CP014743) as a reference for a BLAST search. We identified regions on the WP2202 plasmid that had high percent identity (99.5%) to *C. coli* contigs containing *tetO* from this study. This result suggests that *tetO* is either located on a plasmid or has been integrated into the chromosome, even though no plasmid replicons were identified.

The class D beta-lactamase structural gene, *bla*_OXA-61_, that confers resistance to penams, cephalosporins and carbapenems (Alfredson and Korolik, 2005) was found in all 5 *C. coli* isolates and in 9 *C. jejuni* isolates. The *C. coli* isolates were isolated from house 2 of flock 1 and houses 3 and 4 of flock 2, whereas *C. jejuni* isolates carrying *bla*_OXA-61_ were from houses 1, 2 and 4 of flock 3 **(Table S1**). *bla*_OXA-61_ has been previously identified in both *C. jejuni* and *C. coli* isolates obtained from poultry production as well as humans (Griggs *et al*., 2009; De Vries *et al*., 2018; Gharbi *et al*., 2021). Truncation of the upstream sequence of *bla*_OXA-61_ -35 region has been reported by Alfredson et al (2005) to result in wild-type *C. jejuni* beta-lactam-susceptibility (Alfredson and Korolik, 2005). The upstream conserved sequence of *bla*_OXA-61_ of *C. coli* isolates CC32, CC33 and CC36 was 100% identical to the upstream sequence of *bla*_OXA-61_ harbored on a recombinant plasmid (NCBI: pGU0401) (Alfredson and Korolik, 2005). The other two *C. coli* isolates (CC1 and CC34) had a single T to G mutation at base 66 and CC1 had an additional T to C mutation at base 23. Neither of these mutations were located within the ribosomal binding site or the -10 or -35 promoter regions. We do not know if the presence of *bla*_OXA-61_ conferred the expected resistance phenotype in these isolates since the NARMS *Campylobacter* AST panels used did not include antibiotics classified as penams, cephalosporins or carbapenems. Furthermore, resistance-nodulation-cell division-type multidrug efflux pumps (CmeABC and CmeDEF operons) that confer resistance to antimicrobials and toxic compounds (Akiba *et al*., 2006) were found in *Campylobacter* isolates. CmeABC was present in all *C. jejuni* isolates while CmeDEF was found in both *C. coli* and *C. jejuni* isolates. CmeR is a known transcriptional regulator of CmeABC and when absent or mutated leads to the overexpression of the CmeABC efflux pump and increases levels of resistance to several antimicrobials (Lin *et al*., 2005). While the *cmeR* gene was present in all *C. jejuni* isolates, it was absent from *C. coli* isolates CC32, CC33, CC34 and CC36. In addition to ARGs, we found genes that contribute to arsenic resistance within *C. jejuni* isolates. The arsenical-resistance membrane transporter *acr3,* the putative membrane permease *arsP* and the arsenical pump membrane protein *arsB* were found in all *C. jejuni* isolates from flocks 1 and 2 but absent from flock 3 isolates.

In summary, AST revealed that phenotypic resistance to antibiotics was only present for tetracycline in *C. coli* isolates. Genomic characterization of ARGs discovered the presence of a beta-lactamase gene in flock 2 and 3 isolates that was absent from flock 1 isolates. In addition, two multidrug efflux pumps, one of which was only present in *C. coli* isolates were identified. Therefore, the distribution of ARGs within the *Campylobacter* isolates differed by flock cohort and by *Campylobacter* species.

### Core genome analysis revealed limited genetic diversity among *C. jejuni* isolates

The following *C. jejuni* isolates were found to be identical based on the alignment of their core genome: CJ5, CJ7 and CJ10; CJ4, CJ11, CJ12 and CJ16; CJ14 and CJ23; CJ19 and 28; CJ6, CJ20, CJ21, CJ22 and CJ24; and CJ39, CJ40, CJ41, CJ42 and CJ43. Each set of identical isolates were from the same flock and of the same MLST but not from the same house (**Figure 1**). The high number of identical core genomes suggest that there is limited genetic diversity within the core genomes (genes present in >=95% of isolates) of isolates of the same species or MLST. The core genome for *C. jejuni* isolates consisted of 1116 genes and the core genome for *C. coli* was 1561 genes (**Table S14)**.

Using the Roary-generated core genome alignment, we estimated a maximum likelihood phylogenetic tree using RAxML under a GTR substitution matrix. The resulting phylogeny clustered isolates of the same species or MLST into separate clades. For instance, separate clades were identified for *C. coli* isolates, ST48 *C. jejuni* isolates from flock 3 and *C. jejuni* ST464 isolates from flocks 1 and 2 (**Figure 5**).

**Figure 5.**
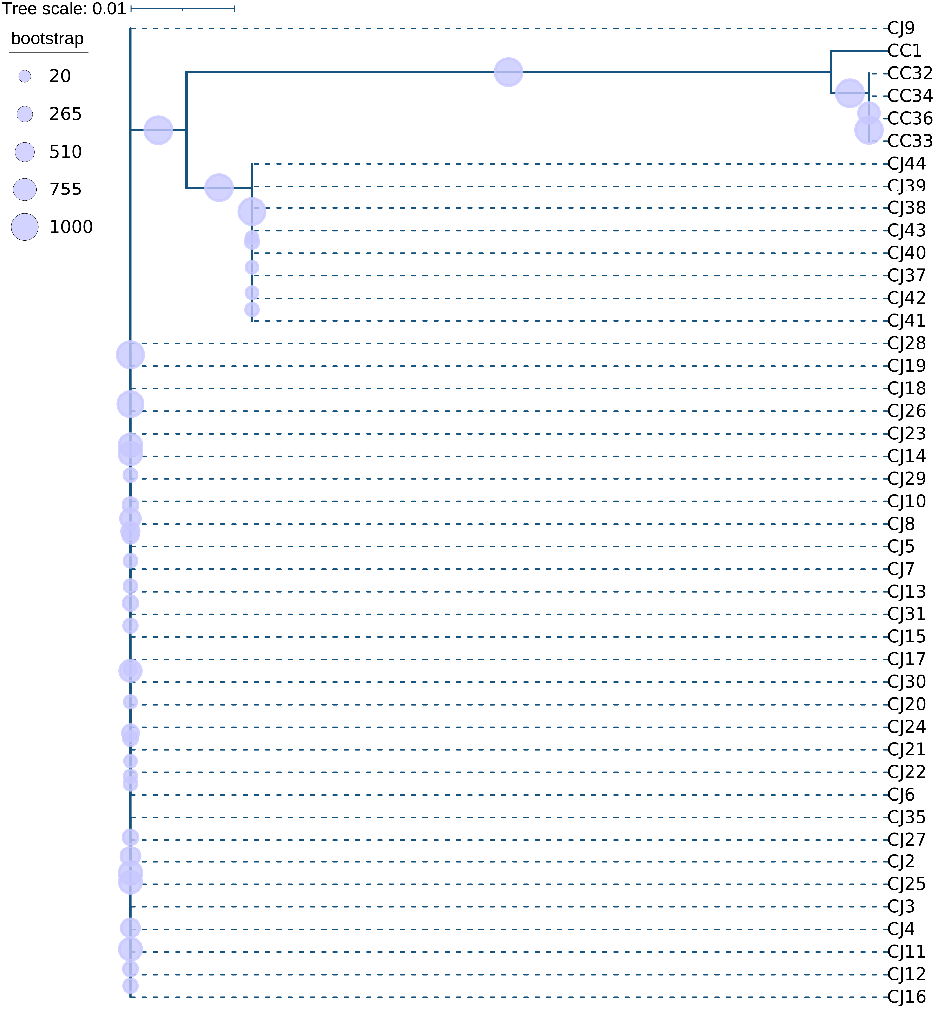
RAxML maximum likelihood phylogenetic tree estimated from the Roary core genome alignment under a GTR model of evolution. 1000 bootstraps were performed to ensure nodal support. Tree was visualized using the Interactive Tree of Life (iTOL).

*C. coli* isolates had a distinct accessory genome (genes present in <95% of the isolates) profile as seen by the Roary-produced accessory genome tree which was plotted alongside the gene presence/absence information **(Figure 6, top).** This consisted of 1,223 genes including 557 associated with hypothetical proteins and 666 annotated genes. Additionally, isolates recovered from flock 3 carried a set of accessory genes lowly present in flock 1 and 2 isolates. This collection of 188 accessory genes consists of 147 associated hypothetical proteins and 41 genes with annotations. Although many isolates shared an identical core genome, all isolates were dissimilar from one another based on their accessory genome. Correspondence analysis based on the presence/absence of accessory genes, VFs and ARGs indicated a separation between flock 3 (ST48) isolates and flock 1 and 2 isolates (ST464) (**Figure 6, bottom**). Thus, both the accessory genome and core genomes grouped *Campylobacter* isolates based on their species and multilocus sequence type.

**Figure 6.**
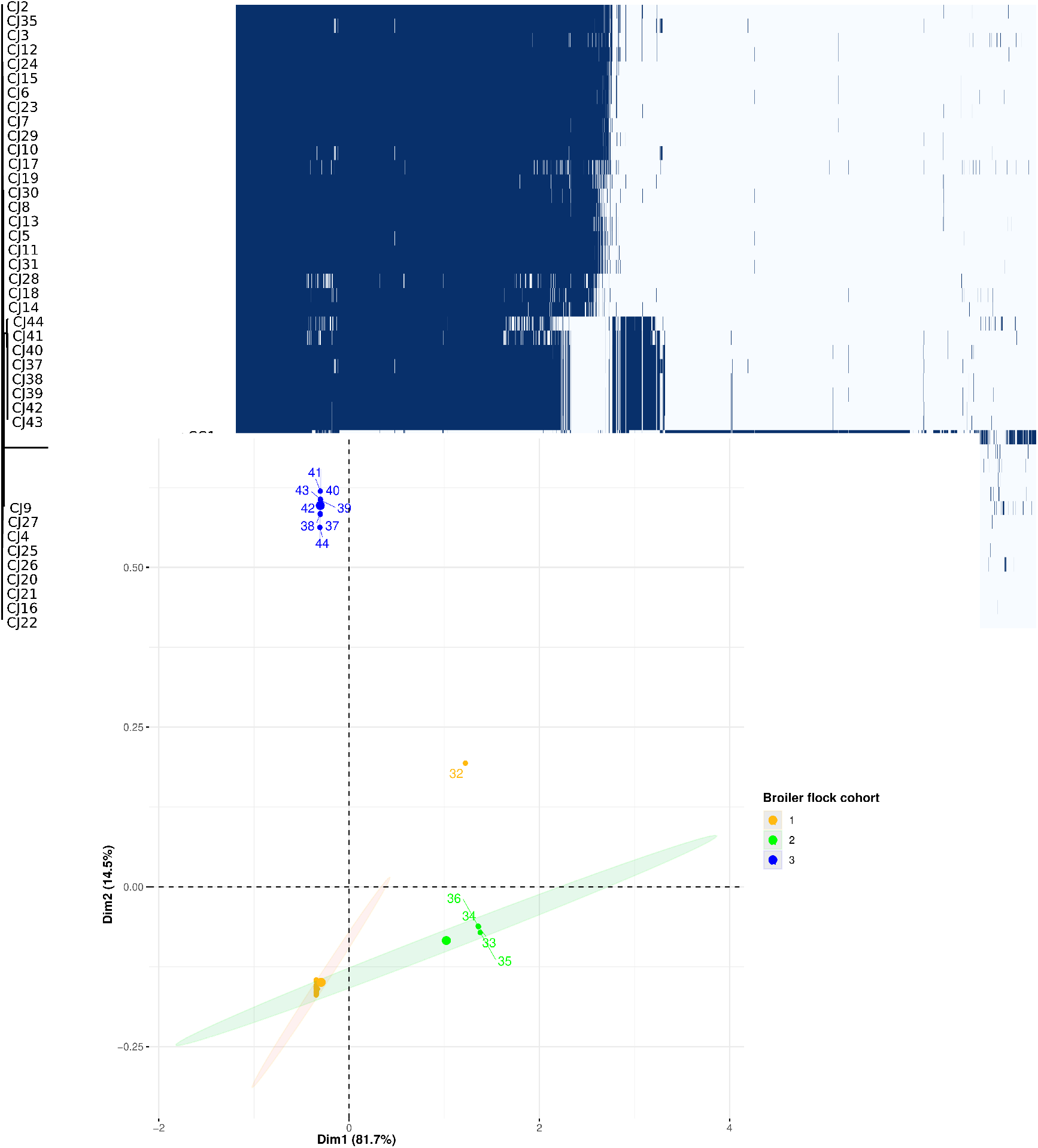
Core and accessory genome analysis. Species labels are denoted with CC (*C. coli*) and CJ (*C. jejuni*). *C. coli* isolates, *C. jejuni* ST48 isolates and *C. jejuni* ST464 isolates harbor distinct accessory genomes. **(Top)** Gene presence/absence matrix of core and accessory genes. The matrix was produced using roary_plots.py and the Roary-generated gene_presence_absence.csv and accessory_binary_genes.fa.newick files. **(Bottom)** Roary, ARG, virulence factor presence/absence correspondence analysis. Correspondence analysis was performed on the combination of the Roary-generated gene_presence_absence.csv and the presence/absence table of ARGs and VFs. Correspondence analysis was conducted in R using factoextra v1.0.7, FactoMineR v2.4 and corrplot v0.2-0 packages.

## Discussion

The purpose of this study was to characterize the ARGs and VFs of *Campylobacter* isolates recovered from litter during 3 consecutive flock cohorts of broiler chickens from 4 co-located broiler houses. Our objective was to identify the ARGs and VFs harbored by these isolates as well as understand how management and environmental factors can lead to genomic changes over the course of multiple flocks.

We found VFs and VF functions that significantly differed across species. VFs relating to adhesion, immune evasion and toxin production differed between *C. jejuni* and *C. coli* (**Table S2**). In general, *C. coli* isolates harbored fewer VF relating to toxin production, adherence, invasion, motility, colonization, and immune evasion than *C. jejuni* (**Figure 4**, **Table 2**). This higher number of VFs in *C. jejuni* may explain why they are more widespread than *C. coli* in broiler production (Powell *et al*., 2012; Whitehouse *et al*., 2018; Tang *et al*., 2020). We also observed differences between *C. jejuni* and *C. coli* in their carriage of T4SS and T6SS. The T4SS aids in the invasion of epithelial cells and has been shown to support intraspecies and interspecies conjugative DNA transfer in *Campylobacter fetus* (Kienesberger *et al*., 2011; Gokulan *et al*., 2013; Van Der Graaf-Van Bloois *et al*., 2016). Similarly, the T6SS is an important VF for *C. jejuni* and it is involved in cell adhesion, cytotoxicity, and invasion *(Lertpiriyapong et al.*, 2012)*. C. jejuni* isolates carrying a T6SS have been identified in poultry and human clinical settings (Bleumink-Pluym *et al*., 2013; Ghatak *et al*., 2017; Kanwal *et al*., 2019; Marasini *et al*., 2020). Kanwal et al. (2019) determined that *C. jejuni* possessing *hcp*, a T6SS gene and important effector protein, had higher hemolytic activity and higher competitive growth advantage against *Helicobacter pullorum*, a bacterium which inhabits a similar physiological niche in chickens (Kanwal *et al*., 2019). No T6SS genes were identified in isolates from the final flock cohort, flock 3, and suggests these genes may impose a fitness cost when in a litter environment. As both the T4SS and T6SS are important virulence factors for the colonization of both chickens and humans, our data suggest that Campylobacter isolates in litter that lack secretion systems will be less likely to infect chickens and therefore less likely to enter into the production facility and consumer-borne food products.

We also found that *C. coli* and *C. jejuni* differed in their susceptibility to antibiotics. *C. coli* isolates (4/5) were resistant to tetracycline while all *C. jejuni* isolates were susceptible to all drugs tested. We identified ARGs and metal resistance genes encoding tetracycline resistance (*tetO)*, arsenic resistance (*arsP* and *acr3)*, multidrug efflux pumps (CmeABC and CmeDEF operons) and class D beta-lactamase structural gene (*bla*_OXA-61_). Tetracycline is both approved for use in food-producing animals and classified as medically important antimicrobial (Center for Veterinary Medicine, 2022). Tetracycline accounts for the largest volume of sales for antimicrobials in food-producing animals and second highest used antibiotic in poultry (Center for Veterinary Medicine, 2022). Thus, it is not surprising we observed tetracycline resistant genes within our isolates. NARMS reporting for Campylobacters does not include beta-lactam/beta-lactamase inhibitor combination agents (Center for Veterinary Medicine, 2022) however, resistance to beta-lactams has been identified in both humans and chickens (Lachance *et al*., 1991; Thwaites and Frost, 1999). For example, *bla*_OXA-61_ has been identified in both human and chicken isolates (Griggs *et al*., 2009; Casagrande Proietti *et al*., 2020).

Species differences were also observed based on the presence of the transcriptional repressor gene for the MDR pump CmeABC, *cmeR*. Mutations, or absence thereof, of the transcriptional repressor, CmeR can lead to enhanced production of the MDR pumps. CmeABC overexpression can lead to reduced susceptibility to tetracycline, ampicillin, cefotaxime, erythromycin, and fusidic acid in *Campylobacter jejuni* (Lin *et al*., 2005) and therefore could explain resistance seen in isolates lacking CmeR. The CmeABC MDR pump and the corresponding regulator, *cmeR*, were present in all *C. jejuni* isolates. *C. coli* isolates that harbored *tetO* harbored CmeABC but not the *cmeR* regulator. The absence of the *cmeR* regulator could have contributed to the level of tetracycline resistance observed in *C. coli* isolates. The presence of this MDR pump, along with its regulator *cmeR*, in the *C. jejuni* isolates suggests that it does not confer resistance, above the epidemiological cutoff value, to the antibiotics tested in our AST panel as all *C. jejuni* were pan susceptible. Overall, *Campylobacter* isolates from this study pose no significant ARG threat and this observation may be attributed to the management program enacted by the producer. Here, the farmer adopted a “No Antibiotics Ever” program after a complete cleanout of the houses were done (Oladeinde *et al*., 2022).

Our previous results (Oladeinde *et al*., 2022) indicated *Campylobacter* was most prevalent during the grow-out of the first flock cohort compared to flock 3 (Oladeinde *et al*., 2022). We observed flock differences with respect to VFs and ARGs. We determined that isolates recovered from the same flock cohort had similar VFs and VF associated functions. Isolates from flock 1 were raised on fresh peanut hull while flocks 2 and 3 were raised on the reused litter from flocks 1 and 2, respectively. We observed a significant difference in VFs in flock 3 (ST48) isolates compared to flock 1 isolates (ST464). We hypothesize that as the litter was reused over multiple flock cohorts the litter microbiome underwent significant flux. The VFs lost over the multiple flock cycles may have imposed a fitness cost resulting in ST464 isolates not being detected past the second flock cohort and ST48 isolates being detected in the final flock cohort, flock 3. The VFs which were absent from flock 3 (ST48) isolates are associated with survival within the chicken gut: capsule biosynthesis and motile phenotype and may not be essential for survival within the peanut hull litter. For example, the *glf* gene, encoding UDP-galactopyranose mutase, is involved in capsule polysaccharide biosynthesis (Poulin *et al*., 2010) and is a known determinant for invasion, serum resistance, adherence, colonization and modulation of host immune responses (Rojas *et al*., 2019). Alternatively, the introduction of ST48 *C. jejuni* isolates into the final flock cohort could have occurred through other means: 1) human or rodent transmission of these strains into the broiler houses, 2) isolation of present isolates was unsuccessful in the previous flock cohorts, or 3) incoming chicks harbored new strains which were then isolated.

We also found that *Campylobacter* prevalence differed between houses. For instance, there was a higher probability of detecting *Campylobacter* in houses 3 and 4 compared to houses 1 and 2 (Oladeinde *et al*., 2022). Observed house differences are unlikely due to differences in day-old chicks because all chicks originated from the same hatchery and were randomly assigned to houses. VFs significantly differed across grow-out houses (**Table 3**, **Table 4**) and this suggests the house environment played an integral role in selecting for these strains. It is possible that these strains were introduced from the hatchery, as a new ST was detected following the introduction of chicks in the final, flock 3, cohort. It is also plausible these strains were residual contamination from the previous flocks and the cleanup performed was not sufficient to remove them. Consequently, upon placement of new chicks, these strains were able to efficiently colonize the naive gastrointestinal tract and spread through houses 3 and 4. Recently, Yi Fan et al. (2022) showed that cleaning broiler houses with water increased activity of the gut microbiota and reduced *Campylobacter* transmission relative to a full disinfection (Fan *et al*., 2022). Therefore, it is possible that the cleaning procedure used had differential effects on the resident bacterial population in each house. For instance, strains from houses 3 and 4 carried VFs that also play a role in organized biofilm formation (*cheA, cheY, cheV and cheW*), which may allow them to adhere to surfaces and persist through cleaning. Additionally, isolates from house 3, which were all from the first flock cohort, harbored a higher proportion of VFs with functions related to immune evasion, glycosylation system and colonization and immune evasion. Flagellin glycosylation has been shown to affect the adherence and invasion of human epithelial cells (Guerry *et al*., 2006). Thus, while these isolates may be better equipped to evade the immune system of the chicken gut and invade epithelial cells, our data suggest these VFs may impose a fitness cost resulting in an inability to persist over multiple flock cohorts.

We have provided new data on the genome characteristics of *C. jejuni* and *C. coli* isolates recovered from the litter of broiler chickens. We demonstrated that the presence of VFs and ARGs varied by species and by flock. While significantly more VFs were present in isolates from house 3, these isolates were not detected in the final flock (flock 3). Additionally, isolates that were found in the litter of flock 3 were missing several VFs that increase an isolate’s ability to colonize and survive within the chicken host including VFs for capsule biosynthesis, motility and the T6SS. Therefore, these data suggest that the house environment and management practices including the initial house cleaning procedure and the reuse of the peanut hull litter over multiple flocks imposed selective pressure on VFs. Nonetheless, there are several limitations of the study which could have biased our interpretation of the results including the small number of flocks, unknown broiler house conditions, as well as the limit of detection of our sampling methodology for *Campylobacter* isolation. Lastly, results from this study are based on one farm and may not be representative of all farms which reuse peanut hull-based litter for broiler chicken production.

## Materials and Methods

Details of methods used for sampling on farms, litter management and bacterial isolation have been described before (Oladeinde *et al*., 2022). We briefly re-describe some of these methods and present others below.

### Study design

Four broiler houses on a farm in Central Georgia, each containing 22,000 to 24,000 broilers per flock, were selected for this study. Three cohorts of broiler flocks were raised in succession in each of the 4 broiler houses between February and August 2018. Before the start of the study a complete litter cleanout was performed in each of the four houses. Before the first broiler flock was introduced fresh peanut hull litter was prepared in each house. Each successive flock, after the first, was raised on the previous flock cohorts’ litter without any cleanout between flock cohorts. During the downtime between flocks the litter was mechanically conditioned by removing the caked portions. Additionally, during the downtime the litter was treated for ammonium control (typically 1 week before sampling) via topical application of a commercial litter acidifier. For the first 14 days of each flock cohort half house brooding was practiced; chicks were only allowed to occupy the front section of the broiler house until after 14 days. Copper sulfate was added to drinking water. All management procedures used are within the scope for routine industry practices.

### Litter sample collection

From February 2018 to August 2018, a total of 288 poultry litter (PL) samples were collected from 4 co-located broiler houses throughout the study. This represents 96 PL sample collections from poultry houses per cohort of broiler flock raised on the same litter. For each broiler cohort, PL samples were collected both early (< 14 days) and late (days 32 - 38) during the grow-out phase at three different sampling times. During each sampling time from each of the four poultry houses, PL samples were collected from four sections: front, mid-front, mid-back and back. From each section, a pool of three litter grabs were collected, bagged litter was transported in a cooler with icepacks until arrival at the laboratory. Litter moisture content was determined for each litter sample by initially weighing 1 g, drying at ∼106°C overnight, and re-weighing to measure dry weight. Moisture content was determined by the difference. Litter pH was obtained by mixing litter (10 g) with 20 ml water, immersing pH probe into mixture, and recording the reading. Poultry house temperature was also collected during each sampling time.

### Bacterial isolation and identification

For *Campylobacter* species detection, appropriate dilutions of the litter mixture were direct plated to Cefex agar (Remel, Lenexa, KS). Plates were incubated in a microaerobic, hydrogen enriched atmosphere (7.5 % H2, 2.5 % O2, 10 % CO2, and 80 % N2) at 42°C for 48 h. Additionally, aliquots of the litter mixture (4 x 50 ul drops) were placed onto a 0.65 µm cellulose acetate filter placed on Cefex agar. Filters were allowed to dry 30 min before being removed and plates were incubated as above. Enrichment was also performed by adding 1 ml of litter mixture to 9 ml bolton’s broth and incubated in a microaerobic atmosphere at 42°C for 48 h before being transferred to Cefex agar and incubated as above. Presumptive positive colonies were selected based on typical cellular morphology and motility using phase contrast microscopy. Isolates were confirmed using the *Campylobacter* BAX® real-time PCR Assay (Hygiena; Wilmington, DE) according to manufacturer’s directions. Twenty-seven unique litter samples were positive for *Campylobacter* and a total of 53 *Campylobacter* isolates were obtained following the different isolation methods described above. For whole genome sequencing, at least one *Campylobacter* positive isolate was selected from the 27 litter samples. Additionally, if there were multiple positive isolates obtained from the same litter sample using the different isolation methods (i.e., direct plating, filter method, or enrichment), the filter and enrichment isolate were chosen over the direct plating isolate. A total of 44 *Campylobacter* positive samples were chosen for whole genome sequencing.

### Antibiotic susceptibility testing

We performed antimicrobial susceptibility testing (AST) on 5 *Campylobacter coli* and 39 *Campylobacter jejuni* isolates recovered from the litter of broiler chicks following the National Antimicrobial Resistance Monitoring System (NARMS) protocol for Gram-negative bacteria. The following antimicrobials were tested: Azithromycin, Ciprofloxacin, Clindamycin, Erythromycin, Florfenicol, Gentamicin, Nalidixic Acid, Telithromycin and Tetracycline. Antimicrobial susceptibility of *Campylobacter* isolates was determined using the Sensititre semi-automated system (Thermo Fisher Scientific, Kansas City, KS) according to manufacturer’s instructions. Briefly, bacterial suspensions equivalent to a 0.5 McFarland suspension were prepared, aliquoted into a CAMPY panel and incubated at 42°C for 24 h under microaerobic conditions. Minimum inhibitory concentrations were determined and categorized as resistant according to breakpoints based on epidemiological cut-off values as used by the National Antimicrobial Resistance Monitoring System (NARMS; https://www.fda.gov/media/108180/download).

### Whole genome sequencing and processing

Illumina short read sequencing was performed on DNA extracted from *Campylobacter* isolates recovered from litter. Libraries were prepared using Nextera XT DNA library preparation kits (Illumina, Inc., San Diego, CA) following the manufacturers protocol. Libraries were sequenced on the MiSeq platform with 250-bp paired end reads. Genome assembly, antimicrobial resistance gene identification, virulence factor identification, plasmid replicon identification, phage region identification and genome annotation were done using Reads2Resistome pipeline v0.0.2(Woyda, Oladeinde and Abdo, 2023). Online ResFinder (Bortolaia *et al*., 2020) for annotation of acquired resistance genes and additional resistance gene identification was performed using the Resistance Gene Identifier v5.1.1 (RIG) and antibacterial biocide and metal resistance genes were identified with BacMet (Pal *et al*., 2014). MLST was determined using the mlst software (Jolley and Maiden, 2010; Seemann, no date), which utilizes the PubMLST website (https://pubmlst.org/). It was required for all reference database hits needed to meet a minimum identity match of 85%. Reference Genbank plasmids AF301164.1, CP044166.1, CP020775.1 and CP014743 were used to determine the exact locations of type 4 and type 6 secretion systems. Verification of genes using Megablast was performed in Geneious Prime version 2022.2.2.

### Statistical analysis

ARGs and virulence database hits from R2R, ResFinder, BacMet and RGI were filtered utilizing >=85% sequence identity to the reference database query and a table was generated based on the presence/absence of identified genes from all isolates. Correspondence analysis on the presence/absence table of ARGs and virulence factors was done in R using factoextra v1.0.7 (Alboukadel Kassambara, Fabian Mundt, 2020), FactoMineR v2.4 (Lê, Josse and Husson, 2008) and corrplot v0.2-0 (Wei, Taiyun and Simko, Viliam, 2021) packages. Heatmaps were generated using the pheatmap (Raivo Kolde, 2019) package in R. A distance matrix was generated using the jaccard metric via the vegdist() function from the vegan v2.6-4 package (Jari Oksanen *et al*., 2022). Optimal number of clusters was identified using the silhouette method implemented by the fviz_nbclust function from the factoextra v1.0.7 package. hclust() from the stats v3.6.2 (R Team, 2015) package was then utilized to perform hierarchical clustering under the ‘average’ (UPGMA) method using the determined optimal number of clusters. All analyses were done in R v4.0.4 (R Team, 2015) utilizing RStudio v1.2.1106 (RStudio Team, 2016).

### Taxonomic classification and phylogenetic analysis

Taxonomic classification of *Campylobcater* isolates using the quality controlled Ilumina short-read sequences was performed with Kraken2 (Wood, Lu and Langmead, 2019, p. 2). Pangenome analysis of annotated assemblies was performed with Roary (Page *et al*., 2015). A phylogenetic tree of the core genome alignment from Roary (core_gene_alighment.aln) was constructed using RAxML with the maximum likelihood method under a GTR model with 1000 bootstraps (Stamatakis, 2006, 2014). The following command was used for phylogentic tree estimation using RAxML: “raxmlHPC -m GTR -p 12345 -s core_gene_alignment.aln -f a -x 12345 -N 1000 -T 48”. Tree visualization was conducted using the Interactive Tree of Life (Letunic and Bork, 2021).

## Data Availability

All raw isolate data, in FASTQ format, is available from NCBI under the accession number: **XYZ**. All other data are available upon request.

## Acknowledgments

We are grateful to Jodie Plumblee Lawrence and Denice Cudnik for their logistical and technical assistance. This work was supported by the USDA Agricultural Research Service (Project Number: 6040-32000-012-000D). R.W. was supported under the National Institutes of Health (Grant number: 5T32GM132057-04 and 5T32GM132057-03). This research was partially supported in part by an appointment to the Agricultural Research Service (ARS) Research Participation Program administered by the Oak Ridge Institute for Science and Education (ORISE) through an interagency agreement between the U.S. Department of Energy (DOE) and the U.S. Department of Agriculture (USDA) (CRIS project number: 60-6040-6-009). ORISE is managed by ORAU under DOE contract number DE-SC0014664. All opinions expressed in this paper are the author’s and do not necessarily reflect the policies and views of USDA, DOE, or ORAU/ORISE. Any mention of products or trade names does not constitute recommendation for use. The authors declare no competing commercial interests in relation to the submitted work. USDA is an equal opportunity provider and employer.

---

**Table S1:** ARG and VF presence absence spreadsheet **(separate file)**

**Table S2.** Virulence factor-associated functions across all *C. jejuni* and *C. coli* isolated. Virulence factor identification was performed with ABRICATE which utilized the Virulence Factor Database (VFDB). For each identified virulence gene, the associated function(s) were enumerated.

| Virulence Factor Function | Count |
| --- | --- |
| Motility and export apparatus | 1623 |
| Colonization and Immune evasion | 779 |
| Glycosylation system | 436 |
| Immune evasion | 340 |
| Toxin | 312 |
| Adherence | 302 |
| Invasion | 295 |
| Chemotaxis and motility | 220 |
| Secretion system | 44 |
| capsule | 44 |

**Table S3.** Comparison of average proportions of virulence factor-associated functions for C. jejuni isolates which significantly differed in proportion between broiler houses 1 and 2 as determined by the Wilcoxon rank-sum test. Virulence genes associated with each function were enumerated for each isolate and a proportion was calculated using the total number of genes in the study population with the given function. Adjusted p value adjustment was performed by the Benjamini-Hochberg false discovery rate correction method. ‘NaN’ values are due to the inability to compute p values due to average proportion values being identical for all isolates.

| **Function** | **adjusted p value** |
| --- | --- |
| Adherence | 1 |
| Colonization and Immune evasion | 1 |
| Capsule | NaN |
| Chemotaxis and motility | NaN |
| Glycosylation system | NaN |
| Immune evasion | NaN |
| Invasion | NaN |
| Motility and export apparatus | NaN |
| Secretion system | NaN |
| Toxin | NaN |

**Table S4.** Comparison of average proportions of virulence factor-associated functions for *C. jejuni* isolates which significantly differed in proportion between broiler houses 2 and 4 as determined by the Wilcoxon rank-sum test. Virulence genes associated with each function were enumerated for each isolate and a proportion was calculated using the total number of genes in the study population with the given function. Adjusted p value adjustment was performed by the Benjamini-Hochberg false discovery rate correction method. ‘NaN’ values are due to the inability to compute p values due to average proportion values being identical for all isolates.

| **Function** | **FDR-BH** |
| --- | --- |
| Immune evasion | 0.16 |
| Adherence | 0.30 |
| Colonization and Immune evasion | 0.37 |
| Glycosylation system | 0.49 |
| Invasion | 1 |
| Motility and export apparatus | 1 |
| Toxin | 1 |
| Capsule | NaN |
| Chemotaxis and motility | NaN |
| Secretion system | NaN |

**Table S5.**
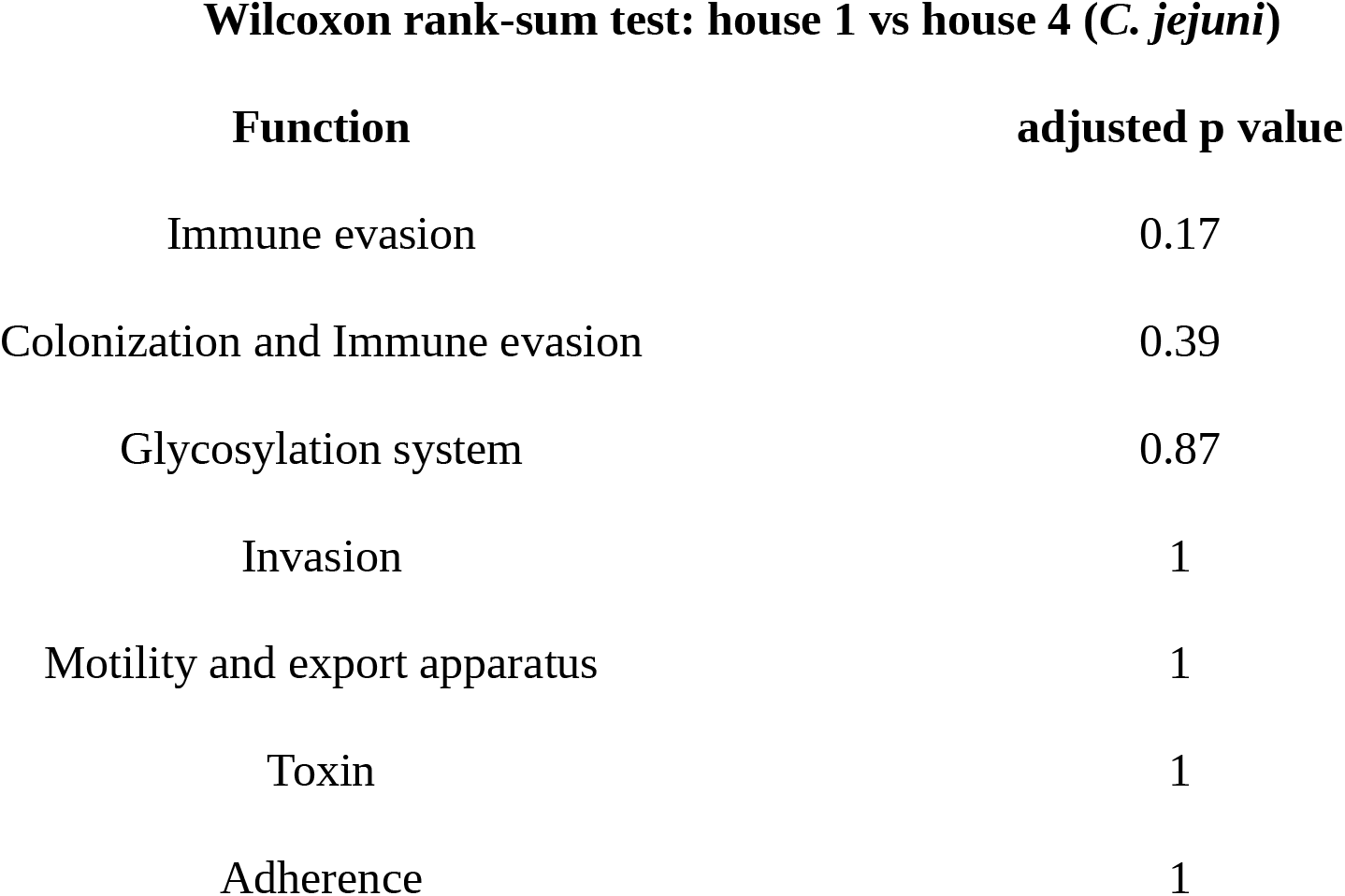
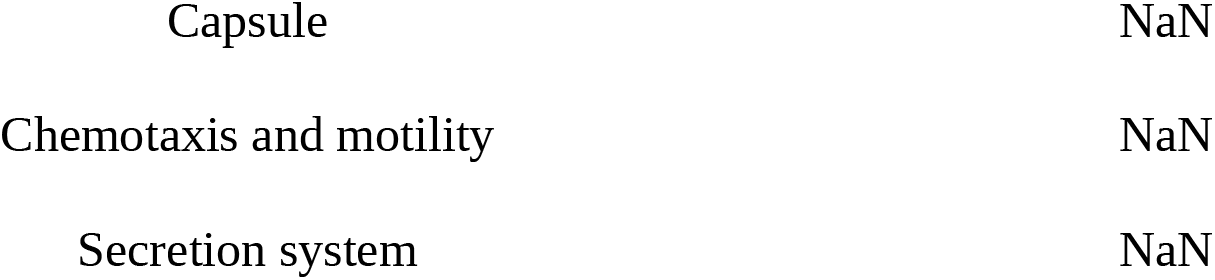
Comparison of average proportions of virulence factor-associated functions for C. jejuni isolates which significantly differed in proportion between broiler houses 1 and 4 as determined by the Wilcoxon rank-sum test. Virulence genes associated with each function were enumerated for each isolate and a proportion was calculated using the total number of genes in the study population with the given function. Adjusted p value adjustment was performed by the Benjamini-Hochberg false discovery rate correction method. ‘NaN’ values are due to the inability to compute p values due to average proportion values being identical for all isolates.

**Table S6.**
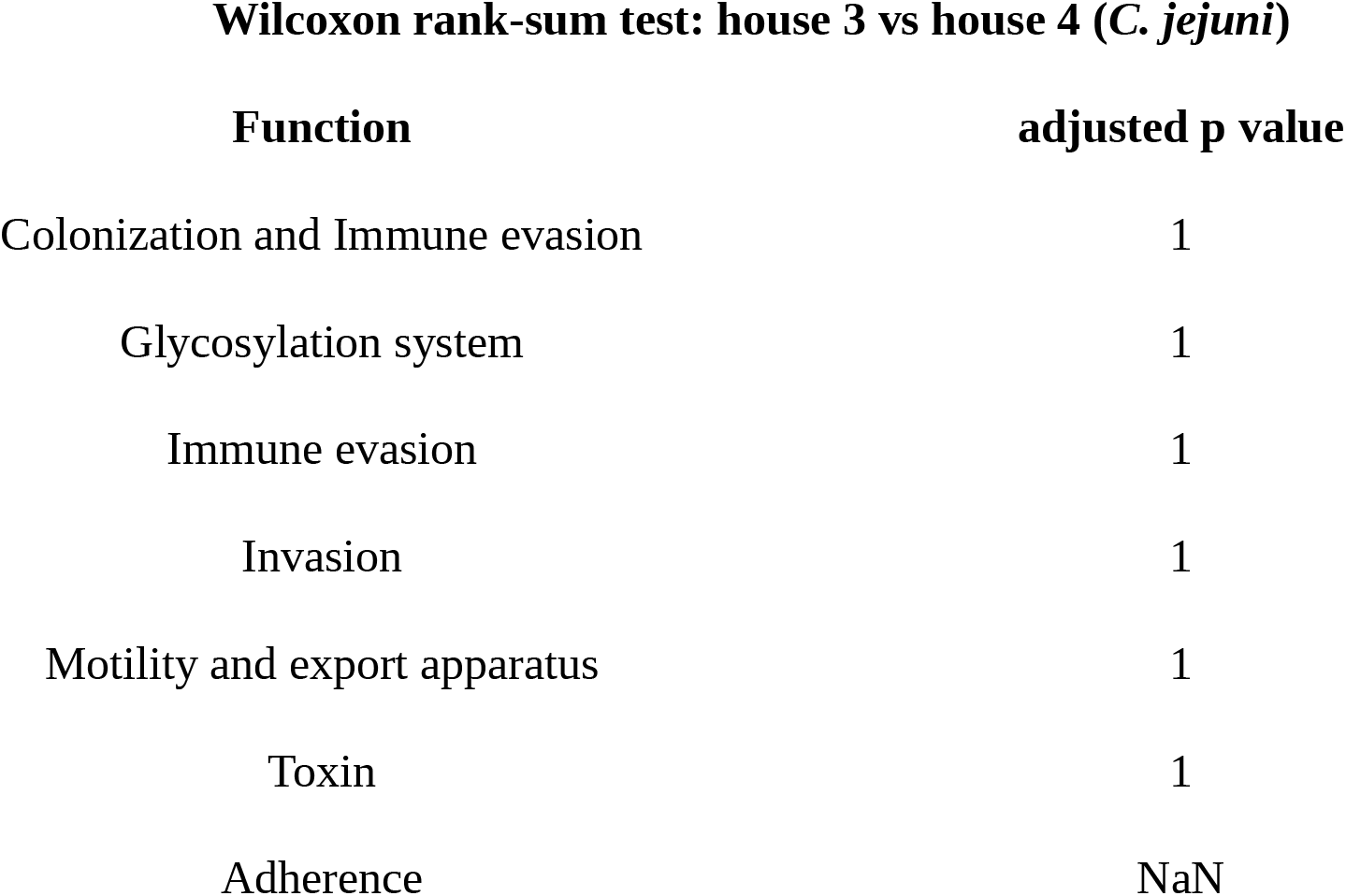
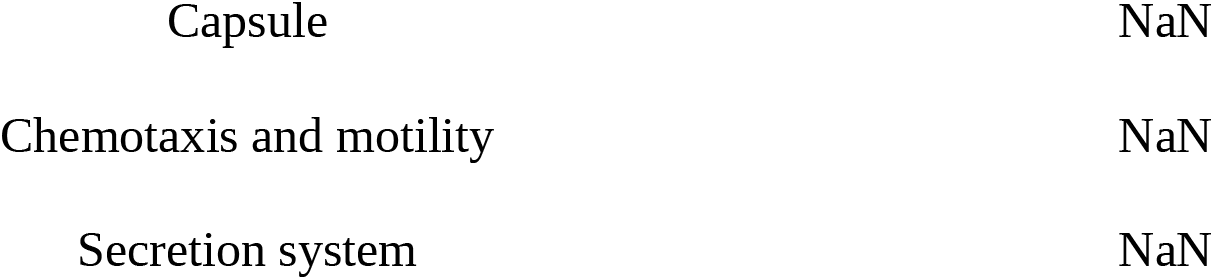
Comparison of average proportions of virulence factor-associated functions for *C. jejuni* isolates which significantly differed in proportion between broiler houses 3 and 4 as determined by the Wilcoxon rank-sum test. Virulence genes associated with each function were enumerated for each isolate and a proportion was calculated using the total number of genes in the study population with the given function. Adjusted p value adjustment was performed by the Benjamini-Hochberg false discovery rate correction method. ‘NaN’ values are due to the inability to compute p values due to average proportion values being identical for all isolates.

**Table S7.** Comparison of virulence factor-associated functions for C. coli isolates from flock 1 and flock 2. Comparison was performed by the Wilcoxon rank-sum test. Virulence genes associated with each function were enumerated for each isolate and a proportion was calculated using the total number of genes in the study population with the given function. Adjusted p value adjustment was performed by the Benjamini-Hochberg false discovery rate correction method. ‘NaN’ values are due to the inability to compute p values due to average proportion values being identical for all isolates.

| **Function** | **adjusted p value** |
| --- | --- |
| Colonization and Immune evasion | 1 |
| Glycosylation system | 1 |
| Immune evasion | 1 |
| Invasion | 1 |
| Toxin | 1 |
| Adherence | NaN |
| Capsule | NaN |
| Chemotaxis and motility | NaN |
| Motility and export apparatus | NaN |
| Secretion system | NaN |

**Table S8.** Comparison of virulence factor-associated functions for *C. jejuni* isolates from flock 2 and flock 3. Comparison was performed by the Wilcoxon rank-sum test. Virulence genes associated with each function were enumerated for each isolate and a proportion was calculated using the total number of genes in the study population with the given function. Adjusted p value adjustment was performed by the Benjamini-Hochberg false discovery rate correction method. ‘NaN’ values are due to the inability to compute p values due to average proportion values being identical for all isolates.

| **Function** | **adjusted p value** |
| --- | --- |
| Immune evasion | 0.63 |
| Colonization and Immune evasion | 0.63 |
| Adherence | 1 |
| Invasion | 1 |
| Glycosylation system | 1 |
| Secretion system | NaN |
| Toxin | NaN |
| Capsule | NaN |
| Motility and export apparatus | NaN |
| Chemotaxis and motility | NaN |

**Table S9:** Virulence factors and functions absent from grow-out cycle 3 isolates.

| Virulence Factor | Function | Function |
| --- | --- | --- |
| ArsP | arsenic resistance | arsenic resistance |
| acr3 | arsenic resistance | arsenic resistance |
| fcl | GDP-L-fucose synthetase | related to capsule - virulence |
| kfiD | UDP-glucose 6-dehydrogenase | related to capsule - virulence |
| PseE.maf5 | Glycosylation system | N-linked protein glycosylation,<br>mutation results in non-motile<br>phenotype |
| glf | UDP-galactopyranose mutase | Galactose metabolism, Amino sugar<br>and nucleotide sugar metabolism |
| Cj1440c | Colonization and Immune evasion | Capsule biosynthesis and transport |
| Cj1432c | Colonization and Immune evasion | Capsule biosynthesis and transport |
| Cj1421c | Colonization and Immune evasion | Capsule biosynthesis and transport |
| Cj1438c* | Colonization and Immune evasion | Capsule biosynthesis and transport |
| Cj1422c* | Colonization and Immune evasion | Capsule biosynthesis and transport |
| Cj1437c* | Colonization and Immune evasion | Capsule biosynthesis and transport |
| Cj1436c* | Colonization and Immune evasion | Capsule biosynthesis and transport |
| Cj1435c* | Colonization and Immune evasion | Capsule biosynthesis and transport |

\* indicates genes absent from grow-out cycle 2

**Table S10.**
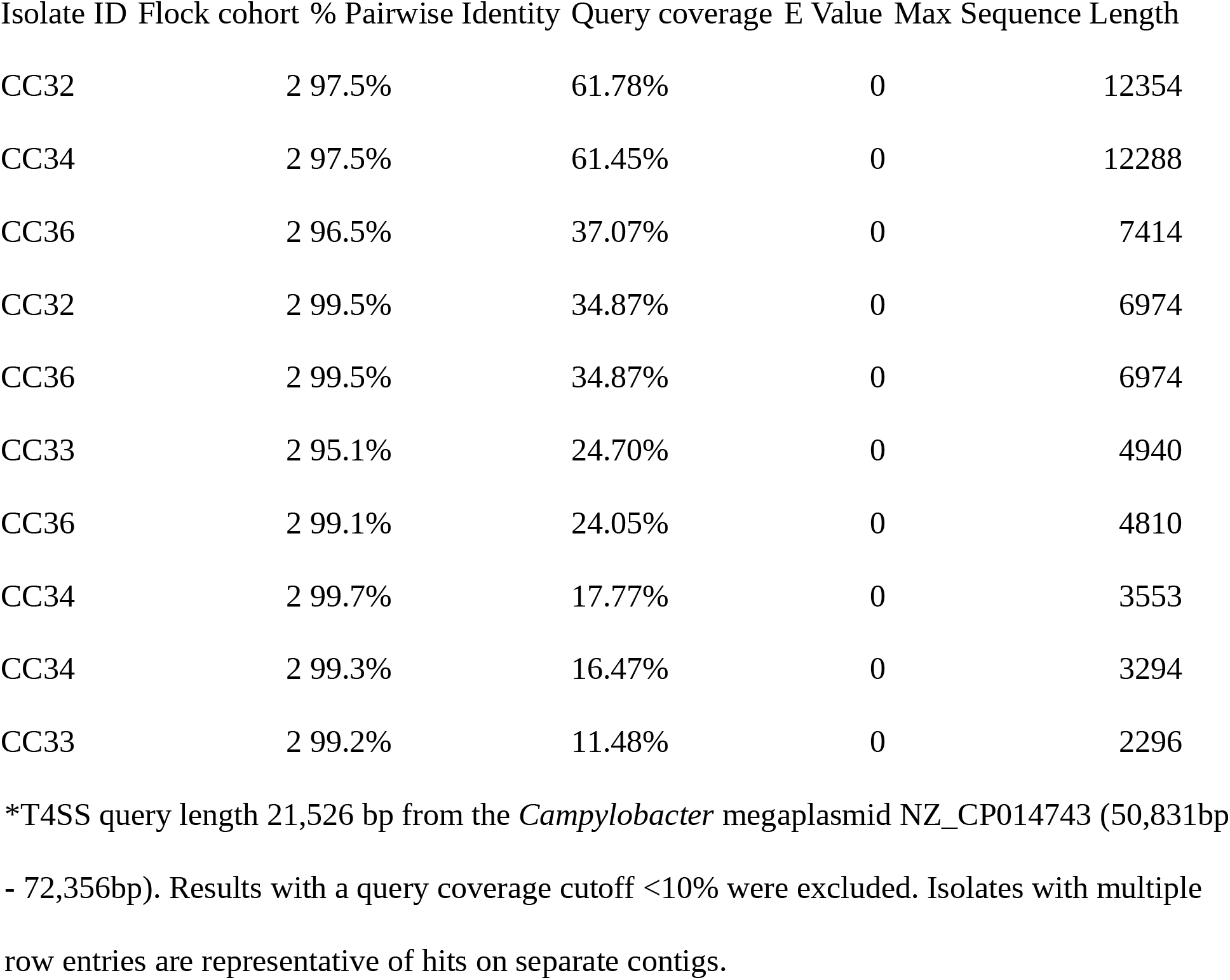
Megablast results for *Campyloabcter coli* isolates against the NZ_CP014743 T4SS.

**Table S11.**
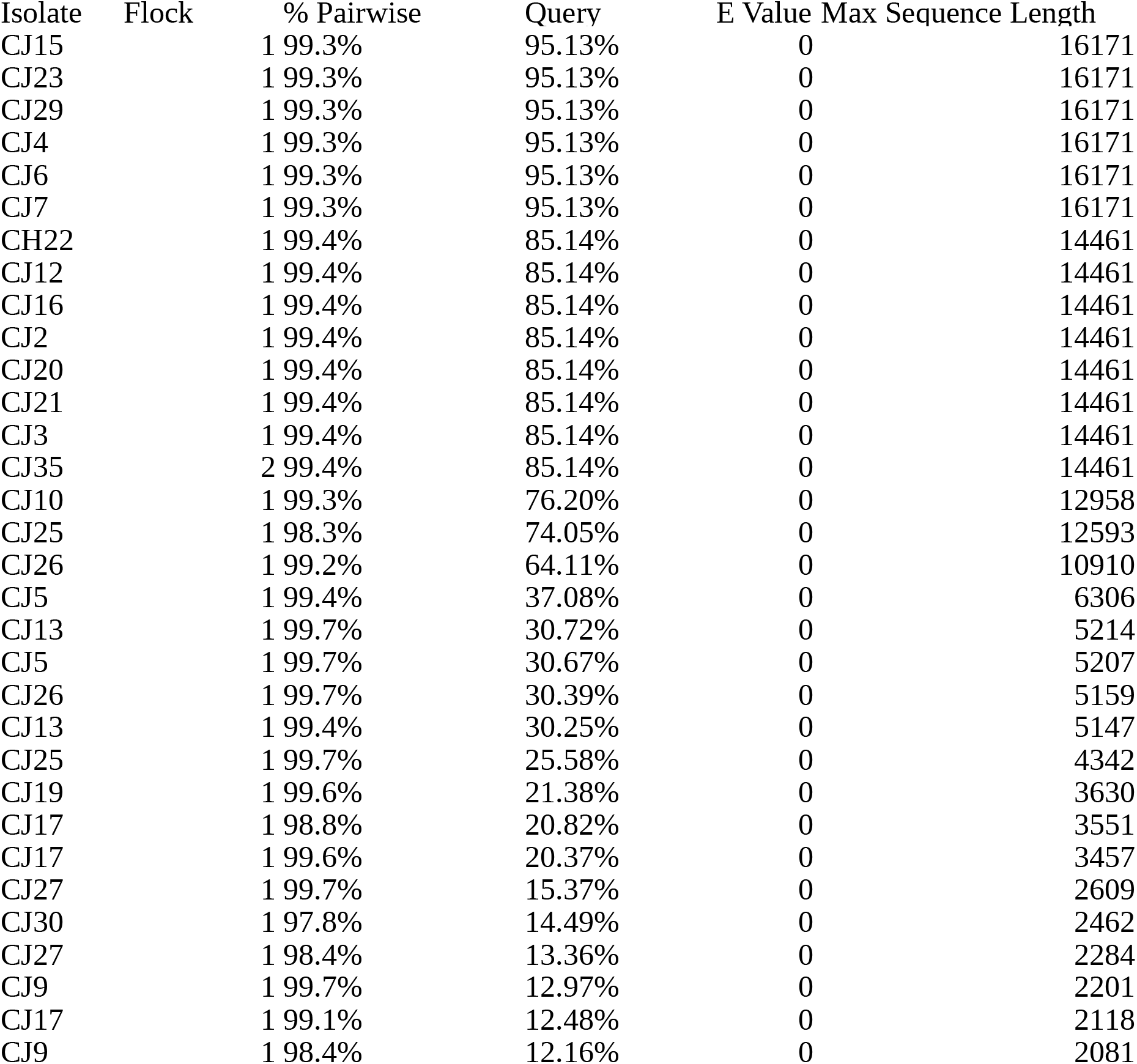
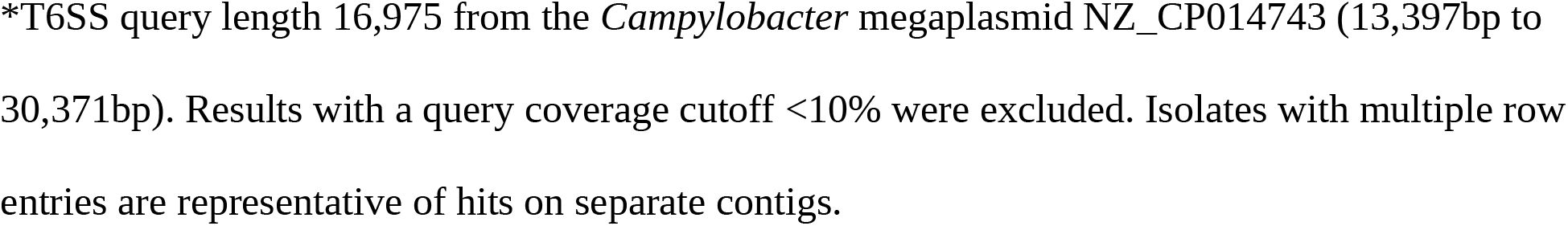
Megablast results for Campyloabcter jejuni isolates against the NZ_CP014743 T6SS.

**Table S12.** Antibiotic susceptibility testing of *Campylobacter* isolates (separate file)

**Table S13.** Antimicrobial resistance genes identified in Campylobacter isolates (separate file)

**Table S14:** Roary presence absence spreadsheet (separate file)

